# Sleep-Dependent Memory Consolidation and Incremental Sentence Comprehension: Computational Dependencies during Language Learning as Revealed by Neuronal Oscillations

**DOI:** 10.1101/218123

**Authors:** Zachariah R. Cross, Mark J. Kohler, Matthias Schlesewsky, M. Gareth Gaskell, Ina Bornkessel-Schlesewsky

## Abstract

We hypothesise a beneficial influence of sleep on the consolidation of the combinatorial mechanisms underlying incremental sentence comprehension. These predictions are grounded in recent work examining the effect of sleep on the consolidation of linguistic information, which demonstrate that sleep-dependent neurophysiological activity consolidates the meaning of novel words and simple grammatical rules. However, the sleep-dependent consolidation of sentence-level combinatorics has not been studied to date. Here, we propose that dissociable aspects of sleep neurophysiology consolidate two different types of combinatory mechanisms in human language: sequence-based (order-sensitive) and dependency-based (order-insensitive) combinatorics. The distinction between the two types of combinatorics is motivated both by cross-linguistic considerations and the neurobiological underpinnings of human language. Unifying this perspective with principles of sleep-dependent memory consolidation, we posit that a function of sleep is to optimise the consolidation of sequence-based knowledge (the *when*) and the establishment of semantic schemas of unordered items (the *what*) that underpin cross-linguistic variations in sentence comprehension. This hypothesis builds on the proposal that sleep is involved in the construction of predictive codes, a unified principle of brain function that supports incremental sentence comprehension. Finally, we discuss neurophysiological measures (EEG/MEG) that could be used to test these claims, such as the quantification of neuronal oscillations, which reflect basic mechanisms of information processing in the brain.

## 1. Introduction

The ability to form memory is essential for an organism to successfully adapt to changing environmental demands (Rasch & Born, 2013). While memory encoding and retrieval occur during periods of wake, sleep facilitates the consolidation of freshly encoded information through unique neuromodulatory activity (Staresina et al., 2015). Electrophysiological research demonstrates that sleep is composed of intensive variations in spatio-temporal oscillations across the brain. These oscillations, characterising rapid-(REM) and non-rapid eye movement (NREM) sleep, originate from signals generated by specific cortical and subcortical networks, and play a key role in memory consolidation (Rauchs et al., 2005).

Evidence suggests the relation between sleep and memory extends to higher-order cognitive domains, such as language (Mirković & Gaskell, 2016; Nieuwenhuis et al., 2013). However, current research on sleep and language is limited to word learning and grammar generalisation (for review, see Rasch, 2017), which does not account for the complex combinatorics of language at the sentence level. Here we propose that sleep is a brain state necessary for the consolidation of the combinatorial mechanisms that underlie cross-linguistic variations in sentence comprehension, namely sequence-based (order-sensitive) and dependency-based (order-insensitive) combinatorics. In addition, we suggest that sleep's effect on the consolidation of sentential combinatorics is reflected in various profiles of brain rhythmicity.

The spatiotemporal architecture of oscillatory rhythms is a fundamental principle of brain structure and function during both wake and sleep states (Buzsaki, 1996; Varela et al., 2001). Sleep-related oscillatory dynamics, such as the sleep-spindle, slow wave oscillation and REM theta activity, will be argued to differentially consolidate sequence-dependent and sequence-independent combinatorics, manifesting in distinct oscillatory activity during sentence comprehension. To support this proposal, we briefly review evidence linking sleep to declarative and procedural memory consolidation, and recent research implicating sleep in language learning. We also review the proposed involvement of the declarative and procedural memory systems in language as posited by Ullman's Declarative/Procedural Model (Ullman, 2001, 2004, 2016). We then outline a new perspective on the involvement of declarative and procedural memory in language by linking mechanisms of sleep-dependent memory consolidation to the neurobiological underpinnings of different types of sentence-level combinatorics (Bornkessel-Schlesewsky & Schlesewsky, 2013; Bornkessel-Schlesewsky et al., 2015). Finally, we will present testable hypotheses arising from this view, focusing on oscillatory brain activity.

## 2. Neurobiology of sleep and memory consolidation

The notion that sleep facilitates memory consolidation and neural plasticity is long-standing (Graves, 1936; Klinzing et al., 2016a). Since its discovery (Aserinsky & Kleitman, 1953), REM sleep was thought to be the sleep stage that supported memory consolidation because of its wake-like EEG and oculomotor activity (Rasch & Born, 2015). However, mixed evidence from studies that selectively deprived subjects of REM sleep (for review see: Vertes & Eastman, 2000) prompted a shift in the sleep and memory field to focus on the role of NREM sleep in memory consolidation. Evidence implicating NREM sleep and associated SWS activity in memory consolidation has given rise to several theories, including the Active System Consolidation (ASC; Born & Wilhelm, 2012; Diekelmann & Born, 2010) and information overlap to abstract (iOtA; Lewis & Durrant, 2011) models, and the Synaptic Homeostasis hypothesis (SHY; Tononi & Cirelli, 2014). According to the ASC model, memory formation is supported by a hippocampal and neocortical system, such that mnemonic representations initially reliant on the hippocampal complex are integrated into the neocortex for long-term storage. From this perspective, sleep integrates hippocampally-dependent memory traces with neocortical long-term memory (LTM) networks by facilitating cross-talk between the two systems (Born & Wilhelm, 2012; Mirković & Gaskell, 2016).

Slow oscillations (SOs; < 1.0 Hz) and sleep spindles (10 - 16 Hz) - hallmarks of NREM sleep - are suggested to be involved in re-processing memory traces within the hippocampo-cortical network (Lewis & Durrant, 2011; Schabus et al., 2004). SOs reflect synchronised membrane potential fluctuations between hyperpolarised up-states and depolarised down-states of neocortical neurons (Klinzing et al., 2016b; Lewis & Durrant, 2011). During phases of depolarisation, sleep spindles are generated from thalamic reticular neurons and promote memory consolidation via cortico-thalamic loops, with individual differences in sleep spindle frequency and density associated with post-sleep memory for motor tasks (Peters et al., 2008), word-pair associations (Schabus et al., 2004), and emotional images (Kaestner et al., 2013). These findings are in line with a broader view (i.e. the ASC model; Born & Wilhelm, 2012) that SOs serve as a temporal gating mechanism for the flow of information between the hippocampus and neocortex, and that the nesting of sleep spindles in phases of depolarisation initiates synaptic change through LTP (Andrillon et al., 2011; Staresina et al., 2015). By contrast, SHY (Tononi & Cirelli, 2014) argues that the plastic processes occurring during wakefulness (e.g. memory encoding) result in a net increase in synaptic weight in networks subserving memory formation. Sleep is argued to facilitate the downscaling of synaptic weight to a baseline level that is homeostatically sustainable; a process posited to be performed by SOs during SWS (Mascetti et al., 2013). This process of synaptic renormalisation desaturates the capacity to encode new information during subsequent wake periods by decreasing neuronal excitability, which in turn, improves the signal-to-noise ratio in the reactivation of stored memory traces (Olcese et al., 2010; Tononi & Cirelli, 2012). The iOtA model (Lewis & Durrant, 2011) builds upon the ASC and SHY models, but makes predictions primarily about schema-conformant memory. According to iOtA, memory traces that are part of the same schemata are preferentially reactivated during sleep via nested spindle and SO activity, and thus develop stronger connections. After synaptic downscaling during SWS, the strongest connections between neurons that share encoded memory traces remain intact, supporting the formation of cognitive schemata (Lewis & Durrant, 2011).

The literature linking sleep and memory consolidation has focused to a large extent on the distinction between declarative and procedural memory, and the unique neurophysiology that contributes to their respective sleep-facilitated consolidation (Smith, 2001). Declarative and procedural memory differ in regard to their level of awareness and the neural networks subserving their computations (Barham, Enticott, Conduit & Lum, 2016). Declarative memory is primarily subserved by prefrontal and medial temporal lobe (MTL) structures, and supports the learning of general facts, namely semantic and episodic memory (Duff & Brown-Schmidt, 2012). In contrast, procedural memory is subserved by a basal ganglia cortico-striatal system, which facilitates the acquisition and execution of motor and sequence learning (Albouy, King, Maquet, & Doyon, 2013; Barnes, Kubota, Hu, Jin, & Graybiel, 2005).

SWS is predominantly associated with the consolidation of declarative memory, assumedly via coordination of widespread neural synchrony that enable interactions between the hippocampus and neocortex (Rasch & Born, 2013). Conversely, REM is assumed to be preferentially associated with the facilitation of procedural memory consolidation, potentially through the activation of locally encoded memory traces in cortical-striatal networks (Barham et al., 2016). It is important to note, however, that the relationship between sleep and procedural memory consolidation is less clear than for declarative memory. In a recent meta-analysis, Rickard and Pan (2015; also see Rickard & Pan, 2017) argue that, for at least finger tapping tasks, sleep does not stabalise procedural memory, and that time of training (e.g. morning/evening), old age (i.e. >59 years), and a build-up of reactive inhibition over training, explain differences in procedural memory consolidation from training to delayed testing over and above that of sleep. There is, however, strong evidence implicating sleep in the consolidation of non-motor procedural tasks, such as auditory statistical learning paradigms (e.g. Durrant et al., 2016), suggesting a beneficial effect of sleep on procedural memory consolidation may be domain-specific.

This claim is corroborated by recent evidence suggesting that EEG phenomena associated with SWS (spindles, slow oscillations) and REM (theta oscillations, increases in acetylcholine; ACh) contribute sequentially to both the consolidation of declarative and procedural memory (Fogel, Smith, & Cote, 2007; Llewellyn & Hobson, 2015). For example, theta rhythms and ACh – which are associated with increased neuroplasticity (Lewis & Durrant, 2011; Llewellyn & Hobson, 2015) – are regulated by REM sleep, such that both increase in the neocortex during REM-rich sleep intervals (Hutchison & Rathore, 2015). These oscillatory and chemical changes – which independently and cumulatively facilitate memory consolidation – support proposals (e.g. the Sequential Hypothesis; Giuditta et al., 1995) that REM strengthens neocortical memory representations that have been selectively refined through the synaptic downscaling of SWS (Cairney et al., 2014; Rasch & Born, 2015; see Figure 1 for a schematic of sleep architecture and associated oscillatory activity in humans).

**Figure 1.**
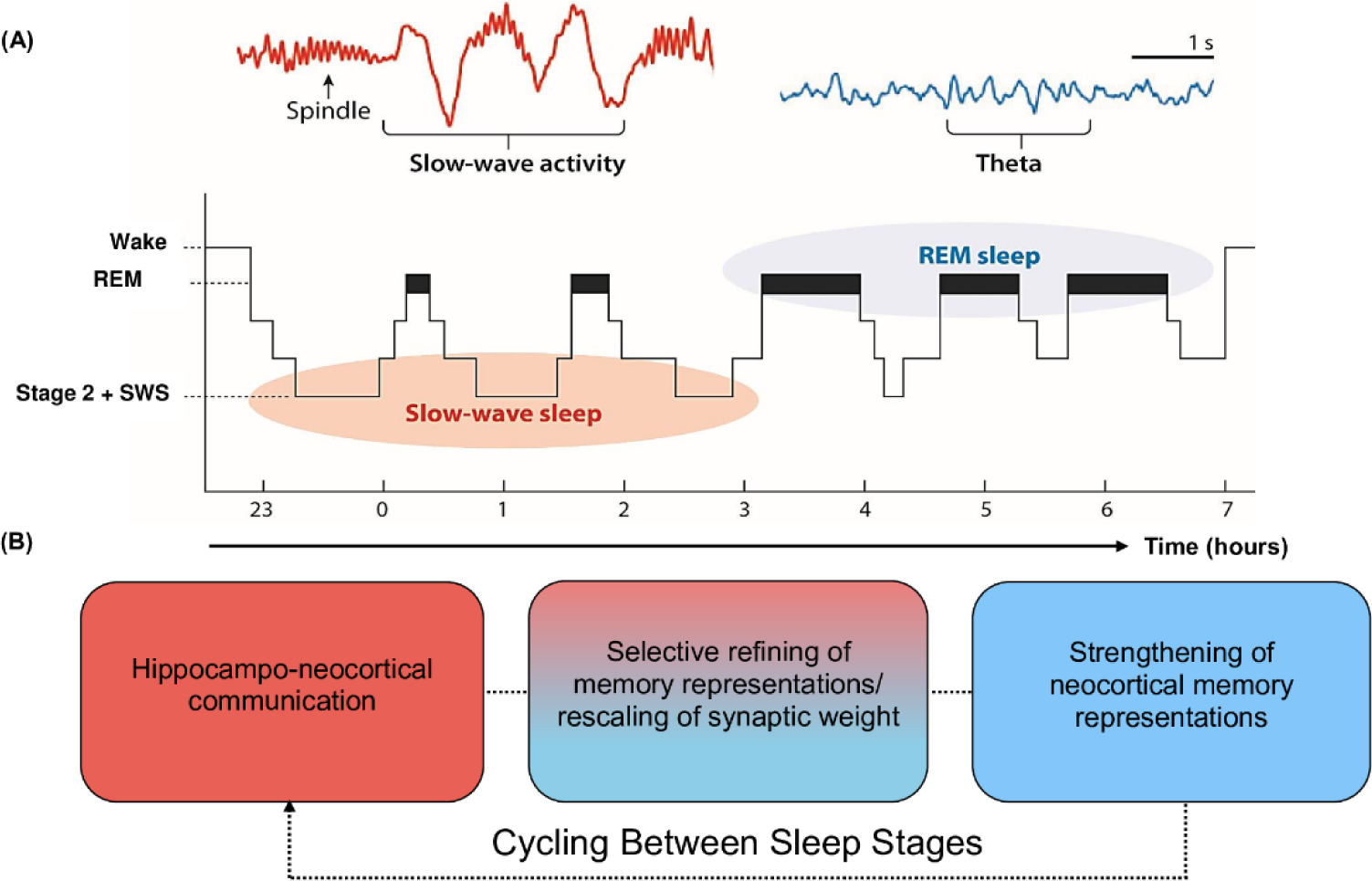
Schematic of sleep architecture in humans and associated oscillatory activity and stages of memory consolidation. **(A)** SWS is most prominent during the first half of the sleep period, and is dominated by neocortical slow oscillations (SOs) and thalamic spindles. By contrast, REM sleep is most prominent during the second half of the sleep period and is characterised by ponto-geniculo-occipital waves, increased acetylcholine (ACh) and cortical theta oscillations (reproduced from Vorster & Born, 2015; permission to reuse image is not required from the copyright holder for non-commercial use as determined by RightsLink®). **(B)** The cyclic occurrence of SWS and REM differentially facilitate memory consolidation. The hierarchical nesting of sharp-wave ripples and spindles within the up state of SOs during SWS facilitate the transfer of information from the hippocampal complex to the neocortex. These neocortically distributed memory representations are strengthened by REM theta oscillations and increases in ACh (Hutchison & Rathore, 2015; Lewis & Durrant, 2011). Each cycle of SWS induces large-scale rescaling of synaptic strength via widespread SO activity.

## 3. A role for sleep in language learning

Interest in the role of sleep during language learning has increased dramatically in recent years, with evidence suggesting that sleep plays a critical role in the consolidation of lexico-semantic information and simple grammatical rules (for review: Rasch, 2017). These experiments consistently demonstrate that sleep consolidates novel word meanings and their respective phonological forms from early childhood by integrating them within the existing mental lexicon (Simon et al., 2017; Dumay & Gaskell, 2007). In particular, SWS promotes novel word production and recognition (Gaskell et al., 2014; Tamminen et al., 2010), and grammar generalisation over and above that of time spent awake (Batterink & Paller, 2015). These findings fit within the ASC model of sleep and memory formation (Born & Wilhelm, 2012). From this perspective, the consolidation of linguistic information occurs during two stages (Davis & Gaskell, 2009; Schreiner & Rasch, 2016).

Initially, the hippocampal complex plays a crucial role in binding a distributed neural representation of the linguistic input, such as word form and meaning (Davis & Gaskell, 2009). During sleep, these newly encoded memory representations are spontaneously reactivated, resulting in localised synaptic downscaling and the distribution of lexical representations in neocortical LTM networks (Rasch & Born, 2013; Schreiner & Rasch, 2016). Thus, sleep is posited to facilitate the integration of newly encoded lexical representations with existing lexical schemata, such as phonological and word form-to-meaning mapping systems (Gaskell et al., 2014).

This idea has been further tested by investigating the effect of sleep on the consolidation of a hidden linguistic rule using event-related potentials (ERPs; Batterink et al., 2014), a derivative of EEG reflecting the synchronised firing of neuronal populations time-locked to specific cognitive or sensory events (Luck, 2014). In this nap study, participants were presented with novel two-word phrases, which included an (English) noun that was preceded by a novel word serving as an article. Unbeknownst to the participants, the novel articles predicted noun animacy, an important semantic feature that is relevant for sentence comprehension in many languages of the world (e.g. Bates et al., 2001). Relative to participants who only experienced SWS, participants who experienced both SWS and REM demonstrated a larger negative ERP occurring between 400-800 ms in response to animacy violations, suggesting greater sensitivity to the hidden linguistic rule. This ERP effect provides preliminary evidence for a modulatory role of SWS and REM in generating neural representations of linguistic information by generalising novel linguistic rules in memory.

This claim is supported by language learning studies that find sleep-mediated effects on oscillatory brain dynamics, suggesting memory-related changes in the organisation of local and distributed neuronal assemblies (Fellner et al., 2013; Hanslmayr et al., 2016; Hanslmayr & Staudigl, 2014). Oscillations within different frequency bands are posited to reflect a number of language-related computations, including the retrieval of newly learned word meanings (Bakker et al., 2015; Takashima et al., 2016) and the detection of violations in artificial languages (de Diego-Balaguer et al., 2011). For example, Bakker and colleugues (2015) reported that novel words encoded before a 12-hr consolidation period elicited greater fronto-temporally distributed theta power at recall than novel words encoded immediately before recall, while de Diego-Balaguer et al. (2011) found that an increase in alpha and theta phase synchrony predicted the detection of violations in learned trisyllabic sequences. Research also reveals that greater theta power during encoding of word-pair associations predicts sleep spindle frequency, which in turn is associated with enhanced recall (Heib et al., 2015; Schreiner et al., 2015; Schreiner & Rasch, 2015). Theta activity is associated with memory encoding and retrieval, and facilitates the consolidation of memory representations in the neocortex via hippocampo-cortical loops (Schreiner & Rasch, 2016), while alpha activity coordinates the flow of information in thalamo-cortical connections that subserve attention and perception (Hanslmayr et al., 2016; Klimesch, 2012; Klimesch et al., 2007). Thus, modulations in theta and alpha activity may differentially modulate the encoding and consolidation of linguistic information, facilitating sleep-dependent memory consolidation, such as spindle-related memory reprocessing (Hanslmayr et al., 2016; Schreiner & Rasch, 2016).

Although this evidence suggests sleep may play a role in aspects of language learning, this does not mean that the consolidation process will be uniform regardless of the material to be learned. Two related factors that may be relevant to memory consolidation, particularly in the case of language, are prior knowledge and systematicity (Gilboa & Marlatte, 2017; Mirković & Gaskell, 2016; Dingemanse et al., 2015).

Prior knowledge has traditionally been viewed as crucial to successful encoding and retention of new knowledge, but research on memory consolidation has revived interest in the notion of schema integration in learning (Gilboa & Marlatte, 2017). A landmark study by Tse et al. (2007) demonstrated that rats’ ability to acquire new associations between flavours and locations depended on the rats' prior knowledge. If new pairings were consistent with previously learned associations involving similar stimuli then the process of consolidation was swift, with new associations quickly becoming independent of the hippocampus. This result is consistent with the idea that a pre-existing mental schema can assist with the learning and integration of new memory traces, a claim supported by McClelland (2013), who demonstrated that this kind of schema-compatibility effect could be explained in the context of a Complementary Learning Systems (CLS) model.

A CLS account predicts that the relationship between the individual elements of a new set of associations can be influential in terms of their initial acquisition and subsequent consolidation. If a set of new associations (e.g., between form and meaning) are in some ways compatible or systematic then they should be acquired more easily by a cortical network with less reliance on the hippocampal complex (Mirković & Gaskell, 2016). If the hippocampus is involved to a lesser extent during initial acquisition, then hippocampo-cortical replay during sleep might also be less important for consolidation, meaning that sleep-facilitated consolidation effects of hippocampally-dependent memory may be weaker. However, such predictions are quite difficult to make because of the potential interaction between prior knowledge and systematicity. That is, the same compatibility in a systematic mapping that leads to weak reliance on the hippocampus during initial acquisition might also lead to greater schema compatibility during consolidation.

Mirković & Gaskell (2016) examined the influence of systematicity empirically in the context of an artificial language learning experiment. They trained participants on a language in which some elements had an entirely arbitrary relationship between the form and the meaning (as is typical of monomorphemic content words), whereas other elements had a more consistent relationship (determiners were used that had a consistent relationship with the gender of the referent). They found that, in this case, only the arbitrary components showed an influence of SWS on performance, consistent with the argument that hippocampal reliance is affected by the level of systematicity.

Nevertheless, several open questions remain. Mirković & Gaskell's (2016) study adopted an afternoon nap paradigm, which occurs at a different circadian phase than nocturnal sleep and is typically dominated by SWS (Payne et al., 2015). In accordance with the sequential hypothesis (Giuditta et al., 1995), interactions between SWS and REM may mediate the influence of prior knowledge and systematicity on the retention of new (linguistic) knowledge. An interactive effect of SWS and REM was demonstrated by Batterink and colleagues (2014), who found that sensitivity to violations of systematic article-noun pairings was predicted by the combined time spent in SWS and REM. From this perspective, during SWS, spindles and SOs may support the consolidation of schema conformant memory (Lewis & Durrant, 2011; Tamminen et al., 2013), while cortical REM theta activity may strengthen systematic mappings between form-to-meaning associations, similar to the beneficial role of REM in facilitating the abstraction of stimuli in probabilistic classification learning paradigms (e.g. Barsky et al., 2015). Thus, while recent research has produced important initial insights on sleep and the consolidation of novel words and simple grammatical rules, we still know relatively little about the neural basis of sleep-facilitated memory consolidation of sentence-level combinatorics, and how an effect of prior knowledge and systematicity may be differentially mediated by different sleep stage characteristics.

### 3.1. Beyond single words: Preliminary evidence for a role of sleep in the consolidation of sentence-level combinatorics

A potential role for sleep in the consolidation of sentence-level combinatorics is identifiable based on studies using artificial and modified miniature languages (MML). Artificial and MMLs generally contain a limited number of words belonging to several syntactic categories that can be combined into meaningful sentences based on the grammatical regularities of a chosen language model (Mueller, 2006). These paradigms provide a useful framework not only to track the learning trajectory of single words, but also the extraction and generalisation of the linguistic building blocks (e.g., sequencing and dependency formation) that underpin sentence comprehension.

Studies using these paradigms (Mueller et al., 2007; Friederici et al., 2002) have helped characterise the neural correlates of language learning by demonstrating that rule violations elicit a biphasic ERP pattern containing a negativity (e.g., N400) and a late positivity (e.g., P600), as observed in natural language studies. Additionally, in a recent functional magnetic resonance imaging (fMRI) study (Weber et al., 2016), speakers of Dutch were exposed to an artificial language made up of thirty-six transitive verbs, ten intransitive verbs and four nouns. Activation in the angular gyrus – a region associated with semantic representations and in unifying smaller concepts into larger representations (Seghier, 2013) – increased linearly across the learning phase (i.e. across 7 - 9 days), and predicted participants’ ability to detect illegal word-order variations. Further neuroanatomical research with shorter learning intervals (i.e. ~ 1-2 days) corroborates Weber and colleagues' findings, demonstrating that hippocampal activation systematically decreases, while activation of language-related neocortical regions (e.g. BA 45 of Broca’s area) systematically increase across (artificial) language exposure (Mueller et al., 2014; Opitz & Friederici, 2003; Opitz & Friederici, 2007). These findings are in line with two-stage models of memory consolidation (e.g. Kumaran et al., 2016; Davis & Gaskell, 2009), further substantiating the notion that newly encoded information is initially reliant on the hippocampal complex before becoming neocortically distributed. As described in Section 2, neocortical LTM networks are strengthened during sleep, suggesting sleep may play a critical role in consolidating language at the sentence-level, but that such an effect may depend on factors related to schema integration and systematicity. Thus, although existing artificial and MML experiments have helped characterise (artificial) language learning, further research is required to expand our understanding of the neurobiological mechanisms underlying the consolidation of language at the sentence-level, such as mechanisms of sleep-dependent memory consolidation.

One model, namely the Declarative/Procedural Model (DP model; Ullman, 2001, 2004, 2016), attempts to ground language processing in the neurobiological systems subserving memory. The DP model argues for a one-to-one mapping between declarative/procedural and semantic/syntactic processing, respectively, and assumes that sleep plays a beneficial role in the consolidation of both memory systems (although it does not provide specific sleep-related predictions; see Ullman, 2016). Since, to the best of our knowledge, the DP model is the only model of language beyond the single word-level that assumes a beneficial role of sleep via the two memory systems as a shared basis, we will briefly review its theoretical underpinnings before introducing our perspective.

## 4. Contributions of the declarative and procedural memory systems to language

For language, differential roles of the declarative and procedural memory systems have been posited and discussed extensively by Ullman (2001, 2004, 2016). It is assumed here that declarative memory underlies the associative memory system required for the mental lexicon and the processing of semantic relations, while procedural memory subserves all rule-based processes in language, including morphology and syntax. As such, the processing of lexico-semantic and syntactic information is argued to differentially engage the neurobiological substrates associated with the declarative and procedural memory systems, respectively.

For the declarative memory system, this is posited to include MTL regions, including the hippocampal complex and entorhinal and perihinal cortices; however, Ullman (2016) recently proposed that there should be a decrease in the involvement of the MTL and an increase in neocortical regions as a function of time and experience of language use. This proposal is in accordance with two-stage models of memory (see Davis & Gaskell 2009 for a discussion on novel word consolidation); however evidence at the sentence-level is limited, and while sleep is assumed to play a beneficial role in the consolidation of both memory systems (see Ullman, 2016), specific sleep-related predictions are absent. By contrast, the procedural memory system is thought to be comprised of parietal, cerebellar, basal ganglia and frontal structures, including premotor regions (Newman et al., 2001; Ullman, 2016). Moreover, Ullman (2001, 2016) argues that specific ERP components are rooted in the neuroanatomical structures of the two memory systems: the N400, which is often associated with lexico-semantic violations (but see, for example, Choudhary et al., 2009; Droge et al., 2015; Frisch & Schlesewsky, 2001; 2005, for evidence against a narrow lexico-semantic function of the N400), is suggested to be tied to MTL and rhinal cortex activation, while left anterior negativities are tied to procedural memory activation (Morgan-Short et al., 2012; Ullman, 2001, 2016). Late positivities, such as the P600, are discussed as originating from ‘conscious syntactic integration’ processes (Ullman, 2016). In regard to (second) language learning, Ullman argues that the declarative system is engaged more strongly than procedural memory during the initial phases of learning, evidenced by greater MTL activation during early second language processing, and greater activation of ganglia cortico-striatal structures when processing becomes more 'native-like'.

Ullman states that "procedural memory should underlie the learning and processing of sequences and rules in language" (Ullman, 2016, p.960), but acknowledges that his predictions for procedural memory are less specific and more tentative than for declarative memory.

## 5. A new perspective on higher-level language combinatorics and the potential role of sleep in their consolidation

The basic assumptions of the Declarative-Procedural Model of language processing are closely tied to the affordances of processing English and languages of a similar type. However, consideration of a broader range of languages calls for a somewhat more complex perspective on the combinatory mechanisms underlying sentence interpretation (e.g. MacWhinney et al., 1984; Bornkessel & Schlesewsky, 2006; Bornkessel-Schlesewsky & Schlesewsky, 2009).

The assignment of thematic roles to noun phrases (NPs) is a case in point. Thematic role assignment allows comprehenders to determine "who is doing what to whom" in the sentence currently being comprehended, and the way in which it occurs is thought to differ between languages (e.g. Bornkessel & Schlesewsky, 2006; Dominey et al., 2009). Native speakers of English typically interpret the first NP encountered as the actor (the active, controlling participant) and the second NP as the undergoer (the affected participant), irrespective of semantic cues (MacWhinney et al., 1984). By contrast, in languages like German, Turkish or Japanese, thematic role assignment is based more strongly on other cues, such as case marking and/or semantic information, including animacy (for a review, see Bates et al., 2001).

In languages of the German/Turkish type, role dependencies can be indicated by case marking, i.e. changes in the morphological form of NPs depending on their role in the current sentence (akin to the difference between subject and object personal pronouns in English, cf. "**I** saw **her**" versus "**She** saw **me**"). Conversely, in languages like English and Dutch, features of animacy and case marking do not influence thematic role assignment. This is evident, for example, in the observation that a sentence such as "The javelin has thrown the athletes" (Hoeks et al., 2004) can only be interpreted as conveying an implausible meaning. Strikingly, this interpretation is reached in spite of the fact that a simple recombination of NPs and verb would yield an interpretation that is congruent with our world knowledge (i.e. the athletes throwing the javelin). That this alternative – plausible – meaning is not accessible to speakers of English and Dutch demonstrates that positional information determines sentence interpretation in these languages (Bornkessel-Schlesewsky et al., 2011; Bornkessel & Schlesewsky, 2006).

These cross-linguistic dissociations in incremental sentence comprehension are captured in proposals that assume distinct combinatory mechanisms in the brain, namely sequence-based and dependency-based (sequence-independent) combinatorics (Bornkessel-Schlesewsky et. al., 2015). From this perspective, speakers of sequence-dependent languages (e.g., English and Dutch) are posited to rely primarily on predictive sequence processing mechanisms for sentence comprehension (Bornkessel-Schlesewsky et al., 2011; Bornkessel-Schlesewsky et al., 2015; Droge et al., 2016). Conversely, speakers of languages such as German and Turkish rely more strongly on sequence-independent features such as case marking or animacy to combine linguistic input into successively more complex representations, thereby facilitating the establishment of relations between non-adjacent elements in a sentence. Note, however, that both types of combinatorics are thought to be operative in all languages: clearly, the processing of languages such as German and Turkish is not completely independent of the order in which the words in a sentence are encountered, and languages such as English allow for non-adjacent dependencies. Thus, rather than being a clear-cut dichotomy, the classification of languages as sequence-dependent or sequence-independent is a matter of degree. This assumed distinction of dependency-and sequencing-based combinatorics as basic and dissociable components of the neurobiology of human language (Bornkessel-Schlesewsky et al., 2015) raises new questions about the relation between these combinatory mechanisms and different memory systems, and accordingly, about the role of sleep in their consolidation. While it appears reasonably straightforward to associate sequence-based combinatorics with the procedural memory system, the status of non-sequence-based combinatorics is less clear. This type of combinatorics is rule-based but sequence-independent (for a similar perspective, see Wilson et al., 2014). It thus shows characteristics of both memory systems (e.g. the requirement for relational binding as in declarative memory; rule-based combinatorics as assumed by Ullman for procedural memory). Consequently, the consolidation of non-sequence-based combinatorics may depend on an interaction between the two memory systems, or may work independently of both systems^1^.

This perspective is closely tied to theoretical advancements in cognitive neuroscience which view the brain as a predictive organ (Friston, 2010; Friston & Buszaki, 2016), and which posit that the (lexico)semantic/syntax distinction can be better described as a segregation of *what* and *when* representations in declarative and procedural memory, respectively. This claim is supported by various neurobiological observations of sleep-dependent memory consolidation -as an optimisation of (Bayesian) model evidence (Hobson & Friston, 2012; Rauss & Born, 2017) -facilitating the generalisation of ordinal sequences (the *when*) and the establishment of semantic schemas of unordered items (the *what*), respectively. These findings provide a promising basis for investigating the consolidation of sequence-dependent and non-sequence-dependent combinatorics from a neurobiological perspective. However, they also demonstrate a need to move beyond the current state of the art in the literature in order to fully capture the complexity of the two types of combinatorics. As described above, non-sequence-based combinatorics involve unordered schemas that are *rule-based* in their organisation; that is, while these schemas are unordered from a sequence-based perspective, they do involve organisational principles of other types. Likewise, sequence-based combinatorics cannot be reduced to ordinal sequences. Rather, they require more richly structured sequence representations, involving asymmetric, hierarchical sequences of elements.

In the following, we derive novel hypotheses about the sleep-dependent consolidation of higher-order language combinatorics based on these assumptions. Specifically, we explore how such hypotheses can be linked to oscillatory brain dynamics, which have long been identified as a key feature of sleep neurophysiology, and which also play an essential role in information processing while awake.

## 6. Sleep-dependent consolidation of higher-order language combinatorics as reflected in oscillatory brain rhythms

Neuronal oscillations are ubiquitous in the central nervous system and play a key role in sensory, motor and cognitive computations during both wake and sleep states (Buzsaki, 1998; Canolty & Knight, 2010). Wake oscillatory activity is typically divided into five bands: delta (δ; ~0.5 --3.5 Hz), theta (θ; ~4 --7.5 Hz), alpha (α; ~8 --12 Hz), beta (β; ~13 --30 Hz) and gamma (γ; >30 Hz; Cole & Voytek, 2017; Mai et al., 2016). Conversely, NREM sleep is predominantly characterised by sigma (12 --15 Hz), δ and slow oscillatory (0 --1 Hz) activity, while REM sleep is dominated by high-intensity, wake-like θ oscillations (Hutchison & Rathore, 2015).

Oscillatory cycles within each band can be conceptualised as temporal receptive windows, transmitting envelopes of information of varying size across or within neuronal pools (Buzsaki & Schomburg, 2015; Harmony, 2013). It follows that slow oscillations, such as those within the δ and θ range, are involved in large-scale network activity, which in turn, modulates faster local events expressed as activity in higher frequencies (e.g. in β and γ activity; Buzsaki & Draguhn, 2004; Sirota et al., 2008). The coupling of activity between fast and slow frequencies allows regions that are part of the same functional network to bind together information that is differentially encoded in memory (Bastiaansen et al., 2012).

Oscillatory neuronal activity is typically quantified using power spectrum analyses, which index local neuronal activity, and phase synchronisation, which is a measure of functional connectivity between distant neuronal populations (Bastiaansen et al., 2012; Rubinov & Sporns, 2010). Possible separable functional roles of each band have been examined across a large body of research in a number of domains, including attention (Klimesch, 2012), memory (Duzel et al., 2010; Hanslmayr et al., 2016) and language (Lewis & Bastiaansen, 2015). Given that oscillatory activity is an inherent property of brain function, supporting both neural plasticity (Hanslmayr et al., 2016) and neural communication (Canolty & Knight, 2010), we posit that neuronal oscillations are a robust means of indexing any effect of sleep on the formation of the neural networks that subserve sentence comprehension. Hypotheses for effects of sleep neurophysiology on oscillatory activity during sentence comprehension are presented below (see Figure 2 for a schematic of the oscillatory mechanisms subserving the encoding, consolidation and retrieval of sentence-level combinatorics).

**Figure 2.**
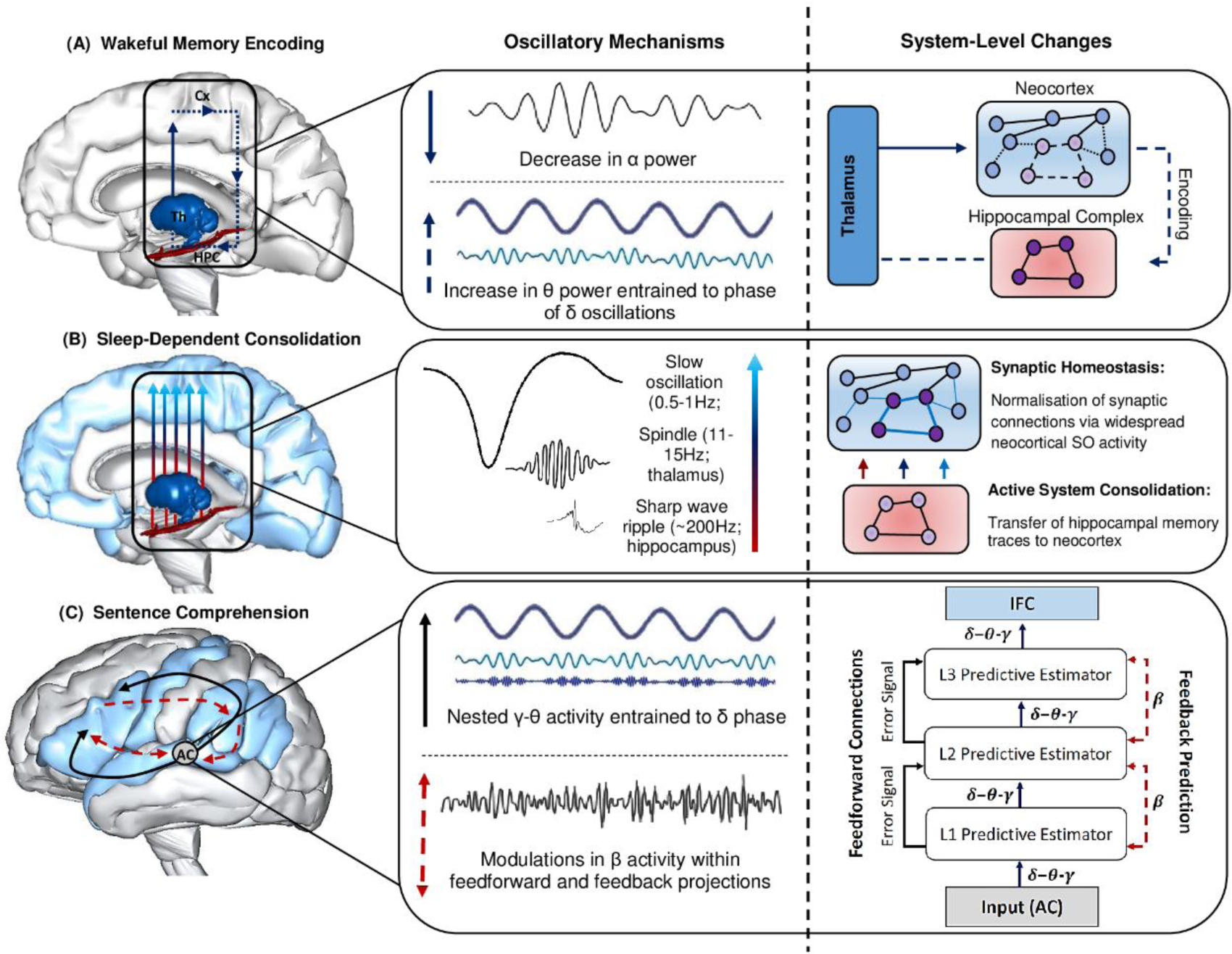
Summary of the oscillatory mechanisms subserving the encoding, consolidation and retrieval of information during language learning and sentence comprehension. **(A)** Decreases in α power facilitate enhanced information processing within the thalamo-neocortical-hippocampal system (TNHs), enabling freshly encoded memory traces to form in the hippocampal complex (HPC). An increase in θ power entrained to the phase of δ oscillations strengthens newly formed memory traces within the TNHs (thalamus, Th; neocortex, Cx). **(B)** Sleep-dependent neurophysiological activity, such as thalamic sleep spindles nested within the up-state of slow oscillations (SOs), enable hippocampal-cortical communication, and the transfer of information to the neocortex for long-term storage, as illustrated by the neural network grids in the right panel. SOs also induce a rescaling of synaptic weight, optimising synaptic efficiency for post-sleep encoding and retrieval. **(C)** The construction of sentence-level meaning is facilitated by long-term memory networks recently established in the neocortex and a fine tuning of synaptic connections in the cortical hierarchy during sleep, as reflected by hierarchically nested δ-θ-γ and β activity, respectively (schematic of hierarchically nested δ-θ-γ oscillations modified from Calderone et al., 2014). Specifically, the schematic in the right panel illustrates the interplay between neuronal oscillations during sentence comprehension from a predictive-coding-based view of the brain following sleep-dependent consolidation. Once acoustic speech patterns are perceived by the auditory cortex (AC), hierarchically nested δ-θ-γ oscillations form successively more complex representations that are generated across the cortical hierarchy (i.e. L1, L2, L3 predictive estimators). Each level of the cortical hierarchy compares feedback (top-down) predictions to lower levels of the hierarchy, a process subserved by β oscillations. Error signals occur when there is a discrepancy between the predicted and actual sensory input, resulting in an update of the internal model. Brain models were generated using BodyParts3D/Anatomography service by DBCLS, Japan.

### 6.1. δ oscillations entrain the activity of higher frequencies

Low frequency oscillations in the δ range have traditionally been associated with memory consolidation processes occurring during sleep (Harmony, 2013; Rasch & Born, 2013). While δ oscillations play a key role in hippocampo-cortical communication during sleep, they also play an active role in sensory processing during wakeful states (Basar & Duzgun, 2016; Schroeder & Lakatos, 2008). Research in the auditory domain indicates that δ-θ oscillations lock to various speech features (e.g. at the syllabic and phoneme level), facilitating the decoding and integration of complex sequences during speech comprehension (Doelling et al., 2014). Similarly, increases in the amplitude of θ and γ oscillations are entrained to the δ phase during attentionally demanding tasks (e.g. oddball tasks), such that increases in θ and γ cross-frequency coupling (CFC) in response to salient stimuli are predicted by the phase of δ oscillations (Calderone et al., 2014; Schroeder & Lakatos, 2008).

In general, the literature suggests that δ oscillations modulate the entrainment of higher frequencies (e.g. θ and γ) during the processing of higher-level information from a task-relevant input stream, such as language and general sequence processing (Canolty & Knight, 2010; Kikuchi et al., 2017). From this perspective, δ oscillations may govern optimal excitability within and between neuronal assemblies, facilitating information transfer during wake, and memory consolidation during sleep. Thus, for language learning, we propose that the phase of δ oscillations will entrain higher frequency bands, facilitating the timing and spiking of synaptic activity, and in turn, facilitating memory encoding. For sentence comprehension, δ activity will depend on the predictability of the category sequence, which will further depend on which units have been successfully consolidated into LTM.

### 6.2. θ oscillations coordinate hippocampo-cortical communication

θ oscillations are generated in the hippocampus and surrounding structures (Covington & Duff, 2016; Piai et al. 2016). They play a key role in the coordination of communication between the hippocampal complex and neocortical regions (Hanslmayr et al., 2016; Herweg et al., 2016). The hippocampus is implicated in relational binding and representational integration, which are important in language processing (Covington & Duff, 2016; Duff & Brown-Schmidt, 2012). Further, the hippocampus has been suggested to be involved in predictive processing by combining elements in memory, supporting the ability to predict future events (Bendor & Spiers, 2016; Friston & Buzsaki, 2016). The process of combining elements in memory to predict sensory input is critical during sentence comprehension, since as information unfolds, the brain generates predictions about upcoming information (Bornkessel-Schlesewsky et al., 2016; Droge et al., 2016). The hippocampus might therefore support language processing by generating predictions for upcoming linguistic information, and θ activity may be modulated depending on whether the sensory input matches the internal model predictions, an idea also recently proposed by Covington and Duff (2016).

This proposal is supported by Friston and Buzsaki (2016, p. 508) who state that “Whether in space or time, ordinal sequences in the hippocampal system may ‘index’ the items (“*what*”) in the neocortex...[and] the organised access to neocortical representations (“*what*”) then becomes episodic [memory] information". From this perspective, during incremental sentence comprehension, the hippocampus may encode the succession of words, accumulating evidence over the duration of the sentence. This evidence may then be used by the neocortex to test *what* predictions about prior beliefs (i.e. the probability distribution of the likelihood of upcoming words based on prior observations). Due to the role of θ oscillations in binding neocortically distributed memory traces (Herweg et al., 2016), θ activity might combine linguistic input into successively more complex representations, establishing relations between (non-adjacent) elements in a sentence. We posit that this may be a general mechanism for the processing of dependencies in linguistic input: dependencies necessarily require the relational binding of two elements; they may also lead to predictions of upcoming input items when the dependent element within a dependency precedes the independent element. From this perspective, dependency processing involves (neocortically computed) relational binding and (hippocampally driven) rule-based processing. Further, as *what* computations are posited to be performed by the neocortex, sleep should optimise the transmission of spatial sequences into more complex (unordered) representations by strengthening connections between the hippocampus and neocortex, and in turn, modulate θ activity.

Modulations in θ power may also index effects of systematicity and prior knowledge on the consolidation of sequence-and dependency-based combinatorics. As discussed in Section 3, newly encoded associations that are compatible or systematic with existing schemata may be acquired more easily by a cortical network with less reliance on the hippocampal complex (Gilboa & Marlatte, 2017; Mirković & Gaskell, 2016; Tse et al., 2007). For example, the consolidation of a second language that shares combinatorial properties similar to a first language (e.g. morphosyntactic case marking in German and Hindi) may result in less hippocampal-dependence during initial learning, and thus weaker sleep-related consolidation effects during SWS (e.g. a reduction in the occurrence of spindles). Rather, effects of sleep may depend on interactions between SWS and REM --as demonstrated by Batterink et al (2014) --resulting in an early emergence of cortical θ activity during incremental sentence comprehension.

In summary, we expect θ oscillations to reveal effects of sleep-facilitated memory consolidation of sentential combinatorics. Specifically, we posit an increase in θ power during incremental sentence comprehension after a language learning task followed by a period of sleep versus an equivalent wake period. We also hypothesise that θ oscillations index hippocampal *when*-based processing, and neocortically-driven *what*-based relational binding, two mechanisms which may depend on hippocampo-cortical communication during SWS and a strengthening of neocortical memory traces during REM. We assume that this effect will accompany both sequence-dependent and sequence-independent combinatory processing, as both types of combinatorics are based on dependency relations. The two types of combinatorics differ in that, on top of basic dependency processing, sequence-based combinatorics include an additional restriction on the positioning of the elements in question as part of a structured sequence.

### 6.3. α oscillations as a thalamo-cortical gating mechanism

According to the inhibition-timing hypothesis (Klimesch et al., 2007), oscillatory α activity modulates the activation of task-relevant cortical regions, facilitating the flow of information through thalamo-cortical networks, and enabling memory traces to form in the hippocampal complex (Bazanova & Vernon, 2014). The generation of α oscillations is posited to occur through GABAergic inter-neurons, an inhibitory neurotransmitter which receives input from excitatory output neurons, manifesting as oscillatory activity in cortico-thalamic and intra-cortical circuits (Mathewson et al., 2011). During NREM sleep, neocortical SOs, thalamic sleep spindles and hippocampal sharpwave ripples facilitate the reactivation of freshly encoded memory traces in the thalamo-neocortical-hippocampal system (TNHs; Bergmann & Staresina, 2017). Thus, α oscillations may serve as a thalamo-cortical gating mechanism, modulating wakeful memory encoding and subsequent reactivation of the TNHs during NREM sleep. From this perspective, α activity might modulate the timing and strength of language learning by facilitating the encoding of novel words and the regularities that govern the combination of words into sentences. Specifically, event-related changes in α power during encoding may index cortical processing in response to novel linguistic information, determining whether sensory input reaches the hippocampal complex via thalamo-cortical connections for long-term consolidation. To this end, we predict that changes in α activity during encoding will modulate language learning outcomes, manifesting behaviourally as greater accuracy of acceptability ratings, and neurophysiologically in distinct oscillatory profiles that reflect successful sentence comprehension. Finally, we expect that this effect will be more pronounced after sleep compared to wake through sleep-dependent reactivation of the TNHs during NREM sleep.

### 6.4. β oscillations reflect a hierarchical predictive coding architecture

β oscillations have recently been proposed to reflect the propagation of top-down predictions to lower levels of the cortical hierarchy during sentence comprehension (Lewis et al., 2016). During highly predictable sentence constructions, β activity is posited to increase, reflecting maintenance of the model predictions. Conversely, β activity is suggested to decrease when prediction errors occur in highly predictable sentences, possibly reflecting mismatches between internal predictions and the actual sensory input (Fontolan et al., 2014; Lewis & Bastiaansen, 2015; Lewis et al., 2015; Weiss & Mueller, 2012). From this perspective, we predict that β power will be modulated by sentences with unpredicted continuations, e.g. sentences deviating from the canonical word order, which are expected to elicit greater beta desynchronisation due to word-order-related prediction errors. We posit that this desynchronisation reflects internal model updates based on mismatches with the actual sensory input, such as the abstract features (e.g. category) and sensory properties (e.g. word form) of the incoming linguistic item (Bornkessel-Schlesewsky & Schlesewsky, 2016). Finally, in addition to the TNHs facilitating offline reactivation of memory traces, homeostatic reductions in synaptic weight during sleep may accentuate prediction error-related β activity relative to an equivalent period of wake (for more on sleep and the formation of predictive codes, see Hobson & Friston, 2012, and Rauss & Born, 2017).

### 6.5. γ oscillations reflect local network activation during the phase of hippocampal θ activity

Our hypotheses for γ oscillations are less specific and more tentative than for the slower frequency bands, since oscillations above ~30 Hz are susceptible to artefact interference, making it difficult to interpret their functional role in information processing and cognition (Buzsaki & Schomburg, 2015; Kovach et al., 2011; Whitham et al., 2007). Specifically, electromyogram and oculomotor signals can contaminate scalp and cortically (i.e., electrocorticography; ECoG) recorded electrical activity >30 Hz, and cause widespread synchronised high frequency oscillations, leading to spurious inter-and intra-regional γ activity (Whitham et al., 2007). Scalp-and cortically-recorded γ activity is also confounded by volume-conduction currents, which result from large fluctuations in subcortical γ rhythms that spread to and inflate γ activity in surrounding cortical layers (for a comprehensive discussion see Buzsaki & Schomburg, 2015). Recordings of cortical neuronal populations are particularly susceptible to volume-conduction currents, as cortical neurons share significant overlap in somatic and dendritic connections (Buzsaki & Schomburg, 2015; Sirota et al., 2008). For this reason, we suggest that the following predictions for γ oscillations be tested with depth electrodes, or at the very least, with magnetoencephalography (MEG), which can overcome spatially spread high frequency activity, since magnetic fields are less distorted by cortical tissue and the low conductivity of the skull (Cuffin & Cohen, 1979; Muthukumaraswamy & Singh, 2013). These approaches would be complemented by advanced analysis techniques, such as independent component analysis, in conjunction with appropriate filtering procedures (Buzsaki & Schomburg, 2015).

In the language comprehension literature, γ synchronisation is argued to reflect accurate model predictions. That is, the matching between top-down (e.g. memory representations of word meaning, contextual information derived from prior discourse) and bottom-up (i.e. the incoming word) information is hypothesised to be reflected in γ synchronisation (Lam et al., 2016; Lewis et al., 2015). However, this research is largely based on cortical (EEG) recordings, which may be confounded by volume conduction currents. Given the possible artefactual nature of scalp-recorded γ oscillations, we will focus on research that has utilised more reliable measures of neurophysiological activity, such as depth electrode recordings.

Research using depth electrodes reveal that γ oscillations occur within the hippocampal complex as well as throughout the cortex (Buzsaki & Schomburg, 2015; Sirota et al., 2008). Further, the selective coupling between regions CA1/CA3 and the medial entorhinal cortex appears to be mediated by γ oscillations that are phase-locked to θ activity (Colgin, 2015). Hippocampally-generated θ oscillations entrain isolated bursts of γ activity through widespread, reciprocal connections between the hippocampal complex and neocortex (Lisman & Jensen, 2013). For example, in a study on waking rats, a large proportion of neocortically generated γ oscillations were dependent on the phase of hippocampally-generated θ oscillations (Sirota et al., 2008). Thus, the temporal organisation between CFC neocortical γ and hippocampal θ oscillations may facilitate information transfer between regionally distant neocortical neural ensembles, which in turn, may support information processing within the hippocampo-cortical system.

This interpretation is in accordance with a θ-γ neural code proposed by Lisman and Jensen (2013), who posit that θ-γ CFC facilitates the generation of ordered multi-item representations within the hippocampo-cortical network, providing information to down-stream regions about the sequence of upcoming sensory input. This interpretation aligns with our proposed role of θ oscillations in dependency-based combinatorial computations, and with Friston and Buszaki’s (2016) perspective on hippocampal *when*-based processing. Within this framework, the hippocampal complex encodes the succession of sensory input, which is then used by the neocortex to perform *what*-based predictions. While θ oscillations support hippocampo-cortical communication, self-organised γ oscillations may help to bind memory representations by (1) allowing neural ensembles that have coded individual memory traces to spike, and (2) generating gaps between temporally encoded items that prevent errors in decoding hippocampally driven sequences, since up to four γ cycles can occur within one θ cycle (Lisman & Jensen, 2013). This proposal is in line with evidence implicating hierarchically nested δ-θ-γ activity in sensory and memory computations, including the perception of speech (see Arnal et al. 2016; Ding et al., 2016; Giraud & Poeppel, 2012). It is also in accordance with the observation that slow cortical oscillations (e.g., δ and θ) reflect large network activation, which in turn modulates the activity of more regionally isolated, faster oscillations (e.g., γ; Sirota et al., 2008).

To this end, bursts of regionally isolated γ activity may reflect the activation of locally encoded memory traces during incremental sentence comprehension, such as the meaning of single words and morphological case marking cues. The entrainment of γ activity to the phase of θ oscillations may then facilitate the binding of these individual memory traces within the hippocampo-cortical network, providing information to down-stream regions about the meaning of the sentence, a process which may be supported by inter-regional δ oscillations. Finally, in line with the notion that SWS and REM play complementary roles in memory consolidation (Giuditta et al., 1995), we posit that REM will strengthen regionally isolated neocortical memory representations that have been selectively refined through the synaptic downscaling of SWS, which will manifest in increased γ activity during incremental sentence comprehension.

## 7. Summary of hypotheses

To summarise, we will restate the above as concrete predictions that follow our proposed functional role of neuronal oscillations in reflecting effects of sleep on the consolidation of sequence-based (order-sensitive) and dependency-based (order-insensitive) combinatorics during language learning and sentence comprehension.

### (a) The phase of δ oscillations entrain the activity of higher frequencies, modifying learning and large-scale neuronal network communication in an attention-dependent manner

Evidence for this prediction stems from research with rodents and monkeys, which demonstrate that δ and θ cross-frequency phase synchronisation coordinates interactions between deep and superficial cortical layers, modifying sensory perception and learning processes, particularly for task-relevant stimuli (Carracedo, 2013; Harmony, 2013). Thus, we hypothesise that the phase of δ oscillations will entrain higher frequency bands, such as θ and γ oscillations, facilitating the timing and spiking of synaptic activity and regulating large-scale network communication during language learning and sentence comprehension. Specifically, δ and θ cross-frequency phase synchronisation will predict enhanced memory consolidation and retrieval, translating into greater accuracy of acceptability ratings during sentence comprehension tasks requiring grammaticality judgements.

### (b) θ oscillations reflect a sleep-dependent transfer of information from the hippocampal complex to the neocortex, and bind relational elements from LTM during sentence comprehension

This hypothesis is supported by intracranial EEG evidence reported by Piai and colleagues (2016), who found that θ power increased in the hippocampal complex during ongoing relational processing during sentence comprehension. In accordance with the general memory literature, θ activity reflects the synchronisation between neocortical regions and the hippocampal complex, binding neocortically distributed memory representations (Herweg et al., 2016; Osipova et al., 2006). This interpretation is supported by sleep and memory research (Bakker et al., 2015; Schreiner et al., 2015), which reports increased neocortical θ power during memory retrieval after a period of sleep, possibly reflecting stronger connectivity between the hippocampal complex and neocortex. These findings are in line with the ASC model (Born & Wilhelm, 2012), which predicts that SOs, spindles and sharp-wave ripples facilitate memory consolidation by modulating hippocampo-cortical communication. Thus, our prediction is two-fold: (1) θ power during incremental sentence comprehension of a newly learned language will be increased following a period of sleep versus an equivalent period of wake, with this increase in power predicted by the occurrence of SOs, spindles and ripples; and, (2) an increase in θ power will occur for both sequence-independent and sequence-dependent interpretation, as both rely on basic dependency formation, which involves the binding of multiple memory traces to form coherent representations.

### (c) Decreases in α power facilitate enhanced information processing within the thalamo-neocortical-hippocampal system, promoting the encoding of novel words and the regularities that govern the combination of words into sentences

α oscillations facilitate cortical processing, acting as a gating mechanism for information flow within thalamocortical loops (Klimesch, 2012; Sadaghiani & Kleinschmidt, 2016). In terms of power, α desynchronisation reflects the activation of cortical areas with increased neuronal excitability (a decrease in amplitude), whereas α synchronisation reflects the inhibition of brain regions (Klimesch, 2012). From this perspective, we hypothesise that α desynchronisation will enhance language learning by enabling novel linguistic information to be processed by the thalamus, promoting the formation of memory traces in the hippocampal complex via the entorhinal cortex. This effect will manifest behaviourally as greater accuracy of acceptability ratings, and neurophysiologically in distinct oscillatory rhythms engaged during sentence comprehension, such as increases in θ-band power during the comprehension of sentences affording a dependency-based interpretation. Finally, we expect that this effect will be more pronounced after sleep compared to wake through sleep-dependent neurophysiology, such that a decrease in α power at encoding and an increase in SOs and thalamic spindles during sleep will predict (1) enhanced behavioural performance on grammaticality judgement tasks and (2) increases in θ-and β-band power during dependency-based and sequence-based processing, respectively.

### (d) During incremental sentence comprehension, β synchronisation reflects maintenance of model predictions, while β desynchronisation reflects prediction error signals

From a predictive-coding-based view of the brain, internal generative models, which predict unfolding linguistic input, update when there is a mismatch between predicted sensory input and the actual sensory input (Bornkessel-Schlesewsky et al., 2015; Pickering & Garrod, 2013). In principle, because of the time-dependent nature of sensory-related predictions, β oscillations may reflect the maintenance of model predictions of general sensory input. From this perspective, during sentence comprehension, feedback projections (reflecting model predictions) that conflict with prediction error signals projected by feedforward connections may increase β-band desynchronisation. This prediction is in line with in *vivo* recordings demonstrating that β oscillations are generated in deep cortical layers, which propagate prediction-related error signals backward on the cortical hierarchy to more superficial layers (Arnal & Giraud, 2012). It is also in accordance with the proposal that β desynchronisation is elicited by bottom-up information that conflicts with top-down predictions during sensory processing (Arnal et al., 2011), or conversely, that β synchronisation occurs when "the cognitive set has to be maintained" (Engel & Fries, 2010, p. 160). Thus, we hypothesise that β power will be modulated by whether incoming linguistic items match internal model predictions. We further posit that SOs will fine tune synaptic connections in the cortical hierarchy, optimising information flow between feedforward and feedback projections, and in turn, optimise accurate model predictions and minimise prediction errors.

### (e) γ oscillations are temporally entrained to the phase of θ and δ oscillations, which subserves the binding of spatially distant neocortical memory traces that have been strengthened during REM sleep

As stated above, our hypotheses for γ oscillations are more tentative than for the slower frequency bands. Based on depth electrode recordings (e.g. Sirota et al., 2008) and MEG research on speech perception (e.g. Ding et al., 2016), we hypothesise that locally generated cortical γ oscillations are temporally entrained to the phase of θ and δ oscillations during incremental sentence comprehension. We further posit that such a hierarchical nesting reflects the following: (1) bursts of regionally isolated γ activity allow neuronal ensembles that code specific memory traces – such as for the meaning of single words – to optimally spike; (2) hippocampally-generated θ activity binds together single memory traces activated by γ activity, and; (3) large-scale δ oscillations facilitate the transfer of γ-θ bound memory representations to regions further downstream. Finally, we predict that increases in REM and associated θ activity after language learning will predict increases in γ synchronisation during subsequent sentence comprehension via a reorganisation of inter-and intracortical memory representations that have been selectively refined during SWS (see Durrant et al., 2015, for a discussion on REM θ oscillations and schema-conformant memory consolidation).

## 8. Concluding remarks and future directions

We have proposed that sleep is an optimal brain-state for consolidating sequence-based (order-sensitive) and dependency-based (order-insensitive) combinatorics. To this end, we argued that sleep-dependent memory consolidation optimises synaptic efficacy, which maximises the ability of the brain to generate predictions of upcoming sensory input during incremental sentence comprehension. We have provided testable predictions for this proposal, focussing on sleep-mediated effects on oscillatory brain activity during language learning and sentence comprehension. δ oscillations entrain the activity of higher frequencies that serve as windows of various size for processing information within and between neuronal pools. α oscillations coordinate the flow of information in a thalamo-neocortical-hippocampal system that subserves memory encoding, and subsequent sleep-dependent memory consolidation. In turn, θ oscillations index a sleep-dependent transfer of information from MTL to neocortex, a process which supports both dependency-and sequence-based combinatorial computations. β oscillations reflect the propagation of predictions and prediction errors via a hierarchically organised predictive coding architecture that is instantiated by sleep-dependent synaptic downscaling. Finally, γ oscillations are entrained to the phase of hippocampally generated θ oscillations, a temporally coordinated process which subserves the binding of spatially distinct, neocortically stored information during sentence comprehension.

Although not within the scope of this paper, it would be worthwhile to consider how mechanisms of sleep-dependent memory consolidation influence the ontogenesis of the functional neuroanatomy of sentence comprehension, such as the dorsal-ventral stream architecture (Bornkessel-Schlesewsky et al., 2015; Brauer et al., 2013). In the visual domain, sleep drives plastic changes in early (V1) and late (i.e. parietal lobe) visual areas, facilitating top-down attentional modulations of primary visual cortex, enhancing visual object recognition (Walker et al., 2005). Similar effects may hold in the auditory domain, such that sleep may trigger large-scale, system-level changes, modifying acoustic memory representations beyond primary auditory cortex, facilitating the recognition of successfully more complex auditory objects (e.g. from syllables to words), a process subserved by the ventral stream (Rauschecker & Scott, 2009; Bornkessel-Schlesewsky et al., 2015).

Clinically, understanding the relationship between sleep neurophysiology and language learning could inform treatments for individuals with language-related disorders, including those with Autism Spectrum Disorder, Specific Language Impairment, and Aphasia, who experience greater sleep disturbances than healthy controls (McGregor & Alper, 2015). Specifically, SOs may serve as a sensitive biomarker of local cortical reorganisation during aphasia therapy post-stroke (Sarasso et al., 2014). Research on both animals and humans indicates that SOs play a homeostatic role in synaptic plasticity by facilitating synaptic depression to obtain a general rescaling of synaptic strength (Tononi & Cirelli, 2014; Sarasso et al., 2014). In this view, if the hypotheses proposed in this paper hold, such that SOs – at least partially – underlie the consolidation of sentential combinatorics, SOs could be selectively increased via stimulation methods (e.g., transcranial magnetic or closed looped stimulation methods; Ngo et al., 2013) to accelerate aphasia-based speech and language therapy. Finally, this paper provides a theoretical framework for understanding how sleep may affect foreign language learning in adults beyond the single word level (e.g. Schreiner & Rasch, 2016), influencing approaches to foreign language learning, which is critical in an increasingly multilingual world.

## Author Contributions

ZC is responsible for the construction and presentation of **Figures 1** and **2**. All authors contributed to the preparation, writing and proof-reading of the manuscript.

## Acknowledgments

We would like to thank Louise Kyriaki, Andrew Corcoran and Alex Chatburn for their valuable comments on an earlier version of this paper, and to Lena Zou for helpful discussion. We would also like to thank Albrecht Vorster, who very kindly provided us with a high resolution image of part A of Figure 1.

## Footnotes

1 Note also that it is not straightforwardly apparent whether existing findings on the sleep-facilitated consolidation of word learning (see Section 3) might generalise to sentence-level combinatorics. The learning of novel words entails the learning of sound sequences, i.e. sequences of phonemes. This raises the question of whether phoneme sequence consolidation operates via similar mechanisms to word sequence consolidation at the sentence level. However, a crucial difference between the two types of sequences is that sequences of words involve the combination of meaningful units into larger meaningful units. Phonemes, by contrast, do not themselves bear meaning, but are rather the smallest units in language that *differentiate* meaning. Consider, for example, the difference between the English words "map" and "nap": here, "m" and "n" lead to a difference in meaning without being meaningful in and of themselves. (See Collier et al., 2014, for an accessible summary of how phonological and syntactic combinatorics differ). It is an open question whether the sequencing property common to both phonological and sequence-based sentence-level combinatorics constitutes a common denominator for mechanisms of consolidation, or whether the two are subject to differential consolidation processes due to the difference in the types of units being combined.

